# Inducible Directed Evolution of Complex Phenotypes in Bacteria

**DOI:** 10.1101/2020.10.30.362871

**Authors:** Ibrahim S. Al’Abri, Daniel J. Haller, Nathan Crook

## Abstract

Directed evolution is a powerful method for engineering biology in the absence of detailed sequence-function relationships. To enable directed evolution of complex phenotypes encoded by multigene pathways, we require large library sizes for DNA sequences >5-10kb in length, elimination of genomic hitchhiker mutations, and decoupling of diversification and screening steps. To meet these challenges, we developed Inducible Directed Evolution (IDE), which uses a temperate bacteriophage to package large plasmids and transfer them to naive cells after intracellular mutagenesis. To demonstrate IDE, we evolved a 5-gene pathway from *Bacillus licheniformis* that accelerates tagatose catabolism in *Escherichia coli*, resulting in clones with 65% shorter lag times during growth on tagatose after only two rounds of evolution.

## Main

Many important phenotypes emerge from the interactions between multiple genes^1,2^. These “complex” phenotypes have traditionally encompassed small molecule biosynthesis^3^, tolerance to inhibitors^4^, and growth in new habitats^5^. However, advances in synthetic biology and metabolic engineering have revealed that even supposedly “simple” phenotypes, such as production of a recombinant protein, become complex as higher performance is desired. This is because auxiliary cellular functions, such as chaperone proteins, cell wall synthesis, and secretion machinery can become limiting in these contexts^6–8^. Clearly, engineering these “systems-level” phenotypes requires systems-level techniques.

In bacteria, complex phenotypes can be accessed via adaptive evolution^5,9,10^ or iterative genome-wide expression perturbation screens (e.g. asRNA^11,12^, and CRISPRi/a^13–15^). One downside of adaptive evolution is the accumulation of genomic hitchhiker mutations, a feature that is particularly troublesome for biosensor-coupled screens and that makes learning from these experiments very time-consuming. For asRNA and CRISPRi/a, the researcher is limited to sampling changes to expression space, rather than the much larger space of protein bioactivity.

For these reasons, directed evolution is useful because it directs mutations to defined DNA sequences and samples a much wider sequence space. However, due to the limited length of DNA that can be evolved using most methods, it has been difficult to apply directed evolution to complex phenotypes. For example, traditional error-prone PCR-based libraries are effectively limited to sequences <10kb in length due to reductions in polymerase processivity, cloning efficiency, and transformation rate above this size. Although recent methods for directed evolution in bacteria have eliminated many hands-on steps via the use of filamentous phage (e.g. phage-assisted continuous and non-continuous evolution (PACE, PANCE)^16–18^, phagemid-assisted continuous evolution (PACEmid)^19^ and Phage-and-Robotics-Assisted Near-Continuous Evolution (PRANCE)^20^), they are limited to small regions of DNA (<5kb) and often couple the mutagenesis and screening steps. This is because the phage used in these techniques (M13) has a strict packaging limit (5 kb) and is engineered to replicate as soon as a certain threshold of biological activity has been reached^21^. Indeed, recent implementations of PACE to evolve multigene pathways (7 kb)^22^ have been limited to relatively small libraries (~10^5^).

Here we add Inducible Directed Evolution, which overcomes these challenges, (IDE, **Figure 1a**) to the directed evolution toolkit. IDE harnesses the large genomes of temperate phages (40-100 kb) to evolve large DNA segments^23^, avoids the accumulation of off-target genomic mutations, and decouples mutagenesis and screening steps. The IDE workflow is both simple and flexible. Pathways of interest are assembled in a phagemid and transformed to a bacterium containing a helper phage. The master regulator for this phage is placed under inducible control. Next, mutagenesis is induced to create random mutations. Then, the phage lytic cycle is induced to initiate phagemid packaging and cell lysis. The resulting phage particles can then be applied to an unmutated strain to start a screening step or another mutagenesis step. We demonstrated IDE using a modified version of the P1 phage (P1kcΔcoi::KanR (ISA137) and P1kcΔcoi (ISA221)) that undergoes lysis in response to the addition of arabinose. Arabinose induces the expression of *coi* to inhibit *c1,* which is the master repressor of lysis^24^. This was achieved by knocking out *coi*, which encodes a repressor of *c1*, and placing it under the control of an arabinose-inducible promoter on a P1 phagemid (PM) ^24^. In the uninduced state, *c1* is expressed and maintains P1 lysogeny. After *coi* induction, *c1* can no longer maintain lysogeny, resulting in P1 particle production and lysis. Inducible mutagenesis was achieved using a previously-described plasmid (MP6)^25^.

**Figure 1.**
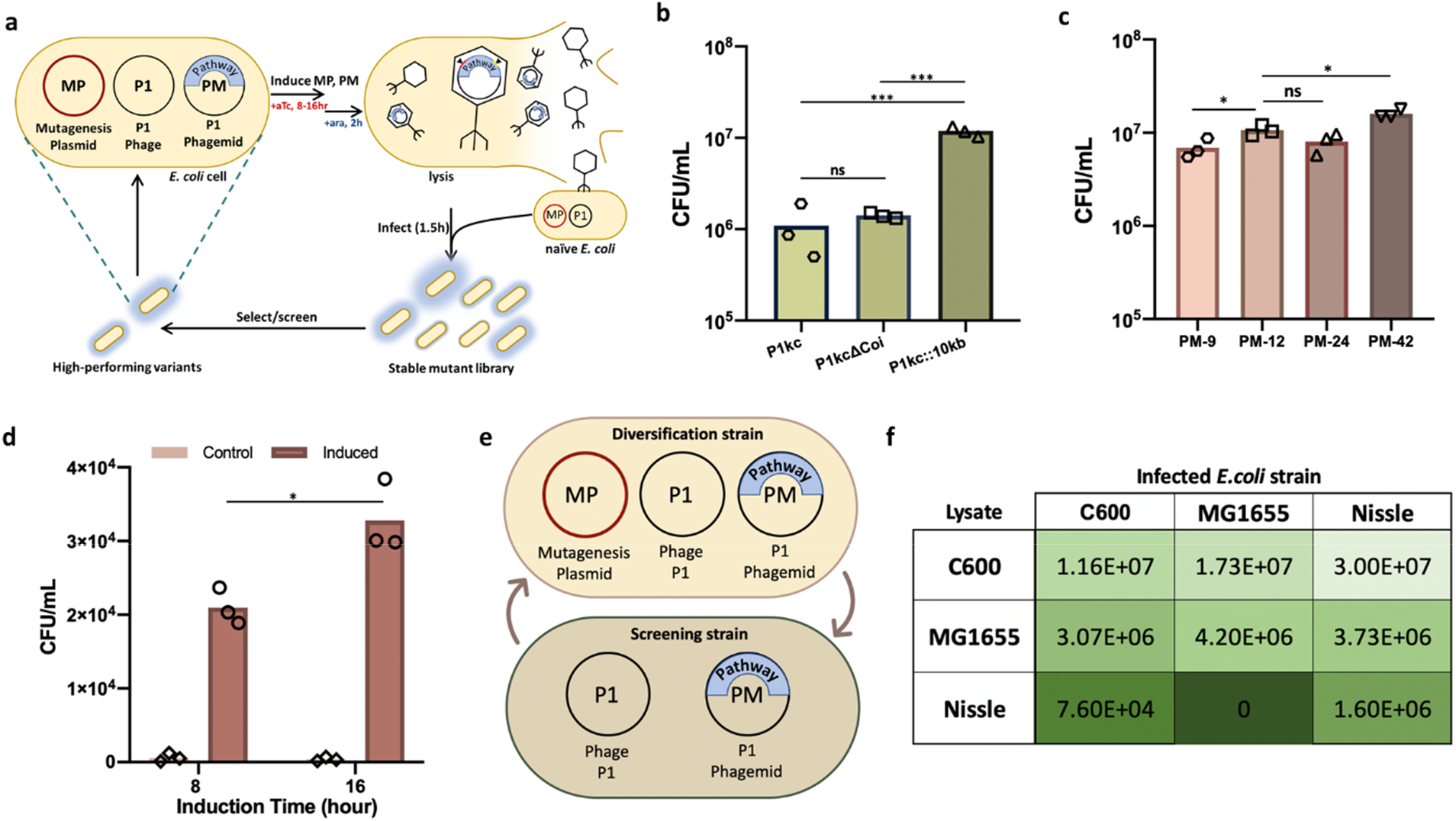
IDE overview and Optimization. **a.** IDE overview. **b.** Engineered P1kc (P1kc::10kb and P1kcΔCoi) increases packaging/infection rates of phagemid compared to wild-type P1kc. **c.** The effect of insert size on phagemid transfer is negligible. PM-9 (9.7 kbp), PM-12 (12.8 kbp), PM-24 (24.0 kbp), and PM-42.0 (42.0 kbp) phagemid were packaged from the same strain (*E. coli* C600) containing P1kc::10kb and same amount of phage lysate is used to infect wild-type *E. coli* C600. **d.** Single stop codon reversion in *CmR*. A single stop codon was introduced to *CmR* and reverted via AIDE, 3 replicate cultures were evolved to gain *CmR* function **e.** Overview of using different *E. coli* strains in an IDE cycle for diversification and screening steps. **f.** Heat map summarizes infection/packaging rates (CFU/mL) of phage lysate produced from different *E .coli* strains (C600, MG1655 or Nissle) and used to infect the same 3 strains.

We first focused on P1 phagemid packaging and infection levels, as these metrics define the number of library members that can be passaged between evolutionary rounds and affect the explorable sequence space. Prior reports of lysate production using P1 were performed by diluting stationary, P1-containing cultures 100-fold into phage lysate medium (PLM) for 1 h at 37 °C followed by induction of lysis via addition of 13 mM arabinose for 1-4 hours at 37 °C^24,26^. The produced phage lysate was used to infect a stationary-phase culture of *E. coli* KL739 at 37 °C for 30 minutes without shaking^24,26^. This lysate production and infection approach produced 2500 CFU/25 μL lysate^24^.

We began by inducing phage production in a much larger *E. coli* C600 culture (OD 1, 5*10^8^cells/mL) containing P1kcΔcoi (ISA221) and PM-12 (ISA012). The resulting lysate (1mL) was applied to 3*10^8^ *E. coli* C600 cells containing P1kcΔcoi and plated on media selective for PM. 6.1×10^5^ PM-containing cells were obtained. To increase this value, we first investigated the composition of the media in which cell growth and infection was performed. Ca^2+^ (CaCl_2_) is required for P1 adsorption to lipopolysaccharide (LPS), and adding Mg^2+^ (MgCl_2_) helps gram-negative bacteria stabilize negatively charged lipopolysaccharides on the membrane^27–29^. Therefore, 100 mM MgCl_2_ and 5 mM CaCl_2_ are commonly added to LB media (forming phage lysate medium (PLM)) in studies of P1 phage^24,26^. We hypothesized that increasing the concentration of MgCl_2_ and CaCl_2_ would enhance P1 infection rates. We found that when the concentration of both salts in PLM is increased by 40%, the number of PM-containing cells increased 2.2-fold to 1.3*10^6^. We called the new medium ePLM (**Figure S1**).

We next varied the optical density to which the recipient cells were grown, while keeping the total number of recipient cells constant. We found that our initial strategy of growing cells to an optical density of 1.0 yielded the highest infection rates (**Figure S2**).

Next, we hypothesized that reducing the ability of P1kcΔcoi to be packaged in phagemids would increase the proportion of particles carrying PM. We found that inserting 10kb of yeast DNA into P1kc (generating P1kc:10kb::KanR (ISA138) and P1kc::10kb (ISA222)) increased transferable library size by an additional 9.2-fold, to 1.2*10^7^ (**Figure 1b,S3**). Similarly, we found that PM packaging was copy-number dependent, with lower PM copy numbers resulting in a reduced number of transduced cells (**Figure S4**). Taken together, over the course of these optimization experiments we were able to increase P1 transduction rates by more than 4,800 fold over previous methods^24,30^.

Although we expected that increasing the amount of lysate applied to naive cells would increase the number of transduced cells, we found that the ratio we had been using (lysate from 5*10^8^ cells applied to 3*10^8^ cells, defined as a ratio of 1:1) was past its saturation level, with reduced amounts of lysate providing similar values (**Figure S3**). Therefore, we varied the number of naive cells, holding the lysate volume constant. As expected, we observed a linear relationship between the number of naive cells and the number of infected cells (**Figure S5**), indicating that IDE library sizes can be easily increased by scaling up lysate and cell amounts.

Using these improved conditions, we investigated the effect of PM size on the number of transduced cells. We observed no reduction in library size with increasing cargo length, up to the largest phagemid we have tested (PM-42, 42.0 kb) (**Figure 1c**). While we expect a substantial reduction of library size with phagemids larger than wild-type P1 (**Figure 1b**), this indicates that IDE is capable of efficiently evolving large multi-gene pathways. In comparison, when these phagemids were transformed via electroporation, the transformation efficiency decreased dramatically as the size of the phagemid increased, up to the size of the P1 genome (90kb) (**Figure S6)**. In addition to the inefficiency of electroporating large plasmids, isolating and transforming large phagemids (e.g. via a kit) is costly and difficult to scale up compared to IDE (simple addition of inducer), making IDE a desirable approach for directed evolution of large phagemids.

Having established efficient transfer of phagemids between cell populations, we next asked whether we could achieve mutations at rates sufficient for directed evolution. For this purpose, we modified a previously-reported plasmid enabling inducible mutagenesis (MP6)^25^. To ensure compatibility with PM, we switched this plasmid to an anhydrotetracycline (aTc)-inducible promoter and a kanamycin selection marker, forming aTc-MP. To test the mutation rate conferred by aTc-MP, we inserted a chloramphenicol resistance gene (*CmR*) with one premature stop codon into PM (ISA308 and ISA311). We expected that inducing aTc-MP would randomly mutate *CmR* and yield variants with the stop codon reverted to a functional codon. We observed time-dependent increases in the number of *CmR*-resistant cells after 16h of induction (up to 5.4*10^−5^ substitutions per base pair (bp), similar to the mutation rate of the original MP625), supporting the notion that IDE enables tunable mutagenesis of defined DNA cargo (**Figure 1d**). Omitting the inducer revealed that aTc-MP has a very tight off state, indicating that aTc-MP is suitable for inclusion in cells during selection or screening steps. Additionally, we found that induction of aTc-MP enabled the simultaneous reversion of two premature stop codons in *CmR* (ISA363) at a rate of 3 per 5*10^8^ induced cells, demonstrating the large library sizes attainable with this mutagenesis technique. The expected reversion mutations were confirmed via sequencing (**Figure S7**).

Because mutagenesis and screening steps are decoupled in IDE, we hypothesized that the strain that is used for screening does not have to be the same as the strain that is used for library generation (**Figure 1e**). This would be beneficial if the ideal screening strain has a limited phage production capacity. As examples, we found that phage lysate produced from 10^8^ *E. coli* C600 cells can passage >10^7^ variants to *E. coli* MG1655 and *E. coli* Nissle 1917 (**Figure 1f**). On the other hand, the same number of *E. coli* MG1655 and *E. coli* Nissle cells can only passage 4.9×10^6^ and 6.9×10^4^ variants back to C600 (**Figure 1g**), respectively. These results indicate that C600 is well-suited for production and packaging of large libraries, enabling these libraries to be screened in a more appropriate strain, for example incorporating biosensors or production-coupled growth circuits.

Because phagemids are transferred to new cells after mutagenesis in IDE, we hypothesized that recessive phenotypes would be easy to select for. Without passage to fresh cells, recessive phenotypes would be difficult to observe due to the presence of multiple plasmid copies. We therefore placed *sfGFP* on PM with medium copy number origin (p15A). Since production of heterologous proteins incurs a fitness cost, cells containing inactivated *sfGFP* would outcompete *sfGFP*-producing cells in a mixed culture^32^. Indeed, after 4 IDE cycles comprising mutagenesis, passage to fresh cells, and cell outgrowth, we found that almost all cells in the culture bore loss-of-function mutations in *sfGFP* (**Figure S8**). The recessive mutations found in this experiment were either premature stop codons or previously-reported mutations that diminish GFP production (**Table S1**)^33^. Loss of GFP fluorescence was not observed in cells that were unmutated (**Figure S8**).

To demonstrate IDE’s capability to evolve a simplistic multi-gene phenotype, we assembled *sfGFP* on a phagemid containing the *pSC101* origin. We chose the *pSC101* origin because of its stringent *Rep101-*dependent replication mechanism and its low copy number^34^. In this setting, GFP fluorescence is controlled by at least 4 different genetic elements (the GFP coding sequence and its promoter, as well as *Rep101* and its promoter). We wished to know which of these elements (or combination thereof) would lead to increased cellular fluorescence when mutated. We found that after two sequential rounds of mutagenesis and passage to fresh cells, (**Figure 2a**), we were able to verify 7 highly fluorescent isolates out of 20 visually selected colonies. Sequencing *Rep101* and *sfGFP* in these 7 isolates yielded mutations exclusively in *Rep101*. To separate these mutations from unknown mutations potentially present in other parts of PM, we cloned these *Rep101* variants into an unmutated PM-*sfGFP* vector and measured fluorescence via flow cytometry. All variants yielded significantly higher fluorescence than wild-type (**Figure 2b**). Most of the *Rep101* mutations present in these clones (R46W, M78I, E93G, E93K, K102E, and E115K) were previously found to increase the copy number of the *pSC101* origin^35^, while one highly beneficial variant (I94N) was novel. It is therefore likely that the increase in GFP production in these isolates is due to an increased phagemid copy number. This result is reasonable because *sfGFP* has already been optimized for high stability and fluorescence in prior studies^36^, and so increasing the copy number of the plasmid may be an easier path to achieve higher GFP expression.

**Figure 2.**
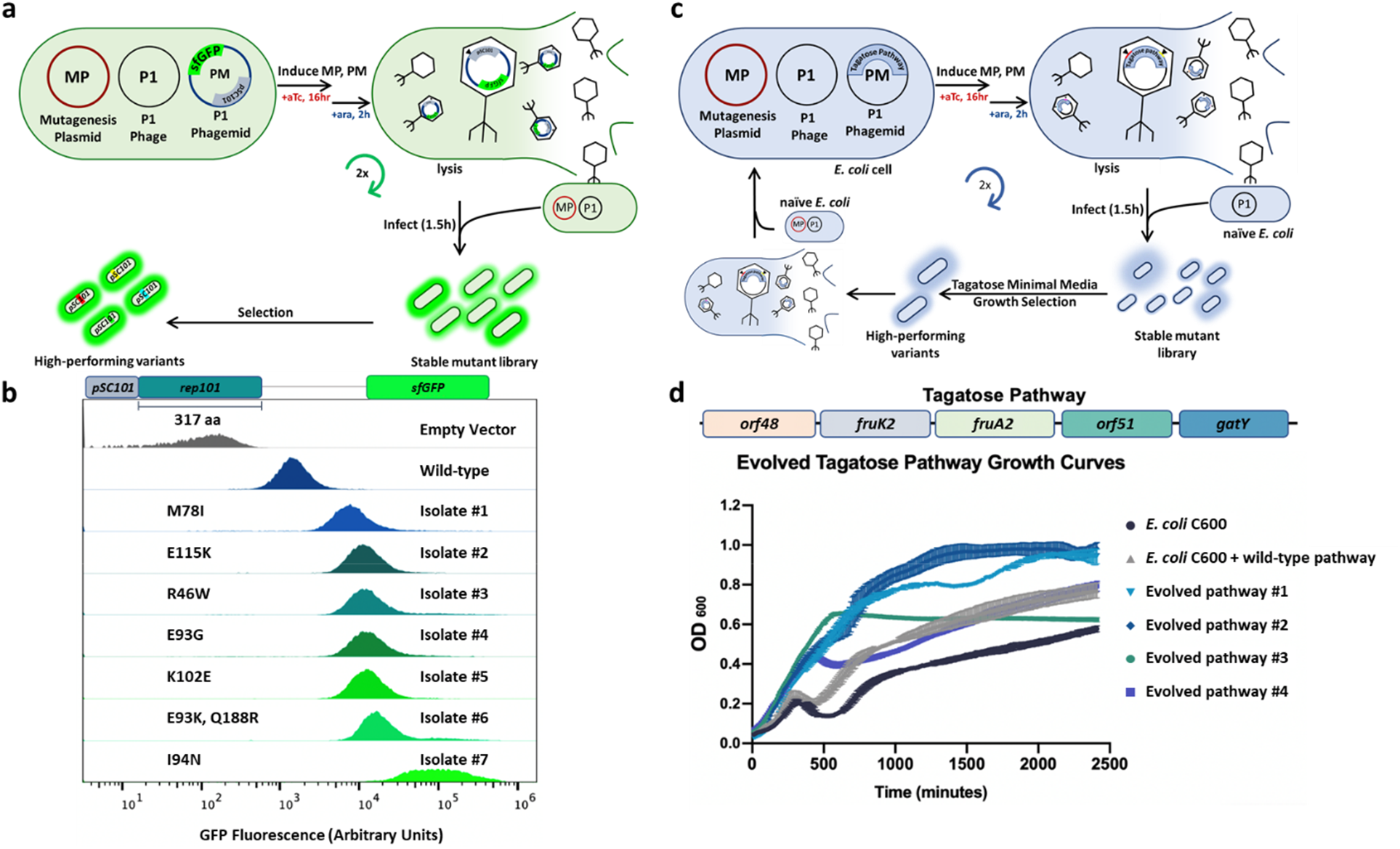
Directed evolution of complex phenotypes. **a.** Overview of evolving pSC101-sfGFP phagemid via IDE. **b.** Flow cytometry histograms of isolated and verified mutants compared to negative and positive controls. All mutations were detected in the *pSC101* origin. **c.** Overview of evolving a tagatose pathway via IDE. **d.** Isolated tagatose pathway variants show improved growth on tagatose minimal media.

Finally, we applied IDE to improve a heterologous tagatose consumption pathway in *E. coli* (**Figure 2c**)*. E. coli* C600 natively consumes tagatose poorly, taking over 12.5 hours to reach an optical density (OD) of 0.25 in media containing tagatose as a sole carbon source from an initial OD of 0.05. We chose a five-gene tagatose consumption pathway from *Bacillus licheniformis* for insertion in PM (**Figure 2d**)^37^. This pathway consists of *orf48* (encoding a predicted transcriptional regulator in the *murR/rpiR* family), *fruA2* and *orf51* (encoding a predicted phosphotransferase system that transports D-tagatose into the cell and converts it to tagatose 1-phosphate), *fruK2* (encoding a predicted kinase that converts tagatose 1-phosphate to tagatose 1,6-bisphosphate), and *gatY* (encoding a predicted aldolase that converts tagatose 1,6-bisphosphate to dihydroxyacetone phosphate and D-glyceraldehyde 3-phosphate). This pathway is therefore a good test case for improving complex phenotypes with IDE, as it encodes different functions that collectively elicit the phenotype of interest. Insertion of this pathway into PM yielded a C600 strain with a 29% reduction in lag time, taking 8.8 hours to achieve an OD of 0.25 in tagatose media (**Figure 2d**). We therefore expected that evolving this pathway would lead *E. coli* to consume tagatose more efficiently. After 2 IDE cycles comprising mutagenesis, growth-based selection, and transfer to fresh cells (**Figure 2c**), we selected for variants that grew faster in liquid tagatose media than strains containing the unmutated pathway. We also performed a parallel selection comprising the same steps, except that mutagenesis was not performed. Phagemids present in cells surviving both selections were transferred to fresh *E. coli* C600, and the growth of strains forming large colonies on tagatose minimal media agar plates was assayed in microtiter plates (**Figures S9 and S10**). Strains from the mutagenic selection exhibited significantly higher growth in tagatose minimal media than strains from the nonmutagenic selection (p<10^-8, Student’s T test) (**Figure S11**). Eight strains from the mutagenic selection exhibiting the best combinations of growth rate and final optical density were cloned into a wild type phagemid backbone (ISA012) to verify increased growth (**Figure S12a**). Of these, four pathways conferred increased growth, relative to the wild-type sequence (**Figure 2c and S12b).** All four variants exhibited some combination of higher optical density (strain E3 exhibited a 2.6-fold higher cell density at 500 minutes than a strain containing the unmutated pathway) and reduced lag time (strain E3 exhibited a 64% reduction in time to reach an optical density of 0.25 than a strain containing the unmutated pathway). Mutations were identified in different genes, as shown in **table S2**. Isolates E1 and E2 have mutations across two sets of 3 different genes (*orf48, fruA2,* and *gatY* for E1, *fruK2, fruA2,* and *gatY* for E2), while isolates E3 and E4 share one mutation in the ribosome binding site (RBS) of *fruK2.* This is the only mutation present in E4, while E3 contains two others (a silent mutation in *fruK2* and a coding mutation in *orf48*) that together further increase growth. The accumulation of fitness-enhancing mutations across multiple genes agrees with prior studies pointing to the utility of a pathway-wide approach to directed evolution^22,38^.

**Table 2.**
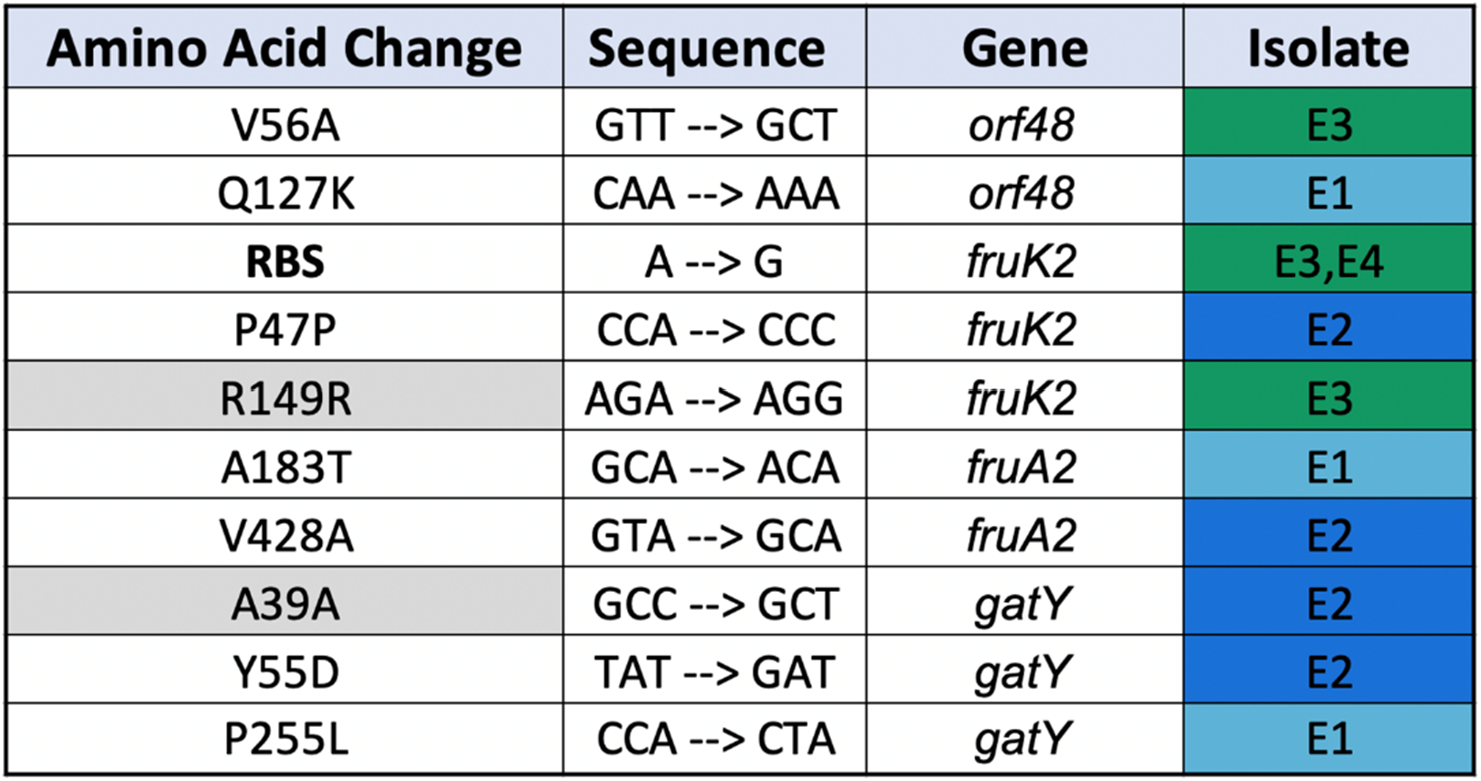
Mutations detected in evolved tagatose consumption pathways. RBS indicates mutations were detected in the ribosome binding site. Grey filling indicates silent mutations. E# indicates the number of the evolved isolate.

As microbial engineering moves toward applications demanding ever-higher performance (e.g. green production of fuels^39^ and chemicals^40^, sensing^41^ and biosynthesis on host-associated sites^42^), the ability to engineer complex phenotypes is becoming increasingly important. Currently, optimizing the performance of multi-gene pathways is a challenging task. IDE offers the ability to perform directed evolution on long (at least up to 42 kbp) sequences of DNA with tunable error rates (up to 5.4*10^−6^ substitutions per bp per generation) and library sizes that scale trivially with culture volume (up to 4% of recipient cells contain mutants). We expect that the use of different mutagenesis methods (e.g. ultraviolet light and chemical mutagens) in place of a mutagenesis plasmid can add further mutational flexibility. Importantly, the use of temperate phages (such as P1) to passage variants to fresh hosts greatly reduces the impact of off-target mutations and decouples mutagenesis and screening steps, providing a large degree of flexibility when designing selections. In particular, we expect this approach to be highly amenable to automation, enabling rapid and highly parallel evolution campaigns similar to the eVOLVER^10^, PACE^18^, and PRANCE^20^ systems. Taken together, IDE demonstrates that temperate phages are promising vehicles for directed evolution of complex phenotypes.

## Methods

### Strains and Media

*E. coli* Top10, NEB 5α and NEB 10β were used for plasmid construction. *E. coli* C600 (CGSC 5394 C600) was used for optimization infection and packaging rates. *E. coli* Nissle 1917 and *E. coli* MG1655 were used to test IDE’s applicability to different *E. coli* strains. All *E. coli* strains were grown in lysogeny broth (LB) (5g/L yeast extract, 10 g/L tryptone, 10 g/L NaCl) at 37 °C supplemented with ampicillin (Amp) (100 μg/mL), kanamycin (Kan) (50 μg/mL) or chloramphenicol (Cm) (34 μg/mL). Strains containing mutagenesis plasmids were grown in LB containing 1% (w/v) d-glucose and appropriate antibiotics. *Bacillus licheniformis* (ATCC® 14580™) was grown in Difco™ Nutrient Broth. All infection and packaging experiments were performed in Phage Lysate Media (PLM, LB with 100 mM MgCl_2_ and 5 mM CaCl_2_) or enhanced PLM (ePLM, LB 140 mM MgCl_2_ and 7 mM CaCl_2_). P1kcΔcoi::KanR and P1kc:10kb::KanR were gifts from Dr. Chase Beisel.

### Preparation and transformation of electrocompetent cells

Overnight cultures of the desired strains were inoculated (1:100 dilution) to 50 mL LB media containing appropriate antibiotics and grown to OD 0.8 at 37 °C and 250 rpm. Cells were then chilled on ice for 15 minutes before being pelleted by centrifugation at 3000 xg for 5 minutes. Cells were then washed twice via resuspension in 25 mL 10% glycerol and pelleting via centrifugation at 3000 xg for 5 minutes. Washed cells were then resuspended in 1 mL 10% glycerol and pelleted at 4000 xg for 3 minutes in eppendorf microcentrifuge tubes. The resulting pellets were then resuspended in 0.5 mL 10% glycerol and divided into 50 μL aliquots that were either used for transformation immediately or stored at −80 °C. Frozen cells were thawed on ice for 10 minutes before being used for transformation. Fresh cells yielded higher transformation efficiency.

### Cloning

Primers used in this study were obtained from Eurofins and are listed in Table S3. Plasmids and phagemids are listed in Table S4. NEBuilder^®^ HiFi DNA Assembly Master Mix was used for plasmid and phagemid construction. SGI-DNA Gibson Assembly^®^ (GA) HiFi 1-Step Kit assembly was used for construction of large phagemids. NEB Q5^®^ Site-Directed Mutagenesis Kit was used to introduce point mutations. Addgene #40782 was used as the backbone for all phagemid cloning, and primers 1 and 2 are used to amplify this backbone for gibson cloning.

### Construction of large phagemids

A 24.0 kbp P1 phagemid (Addgene #40784) was used as a backbone for constructing a 42 kbp phagemid (PM-42.0). To construct this phagemid, 3 PCR reactions were used to amplify 5-7 kbp fragments from the *Saccharomyces cerevisiae* genome in addition to the phagemid backbone. Q5^®^ High-Fidelity 2X Master Mix was used to amplify the parts and SGI-DNA Gibson Assembly^®^ (GA) HiFi 1-Step Kit was used to assemble the parts. The assembled product was transformed into *E. coli* DH10B (ISA585). ZymoPURE™ II Plasmid Midiprep Kit (Catalog No. D4200 & D4201) was used to extract PM-42.0 from ISA585.

### Phage Production

Overnight cultures of strains containing P1 and phagemid were subinoculated in ePLM (1:100 dilution) with appropriate antibiotics. At OD 0.8-1.0 cell cultures were induced with 20% L-arabinose (1/100 culture volume) and put back to the shaking incubator (37 °C, 250 rpm). After 2 hours, the cultures were removed from the incubator and transferred to 15 mL centrifuge tubes containing chloroform. The tubes were left on ice for 5 minutes with gentle mixing or pipetting every minute. The tubes were then centrifuged at 3000 xg for 10 minutes at 4 °C. The produced phage (present in the supernatant) were then transferred to sterile tubes for storage. Phage lysate is stable at 4 °C for 1 year and indefinitely at −80 °C.

### Phage Infection

An overnight culture grown in LB with the appropriate antibiotics was subinoculated into ePLM (1:100 dilution). At OD 1, the cells were spun down at 3000 xg for 5 minutes. The supernatant was discarded and the pellet was resuspended in ⅓ volume fresh ePLM. The cells were then added to phage lysate in a 14 mL falcon culture tube. The infection mixture was incubated in a 37 °C incubator (250 rpm shaking) for 20 minutes and then moved to a 37 °C standing incubator for 20 minutes. The infection mixture was quenched with 1 mL of Super Optimal Broth (SOC) containing 200 mM sodium citrate. The mixture was then incubated for 40 minutes in a 37 °C shaking incubator before being transferred to 50 mL LB media with the appropriate antibiotics or plated on LB agar plates containing the appropriate antibiotics.

### Cell growth phase experiments

Three *E. coli* C600 colonies grown overnight in LB containing the appropriate antibiotics were subinoculated (1:100, 1:200 and 1:300 dilution) into three 50 mL PLM and grown to OD 0.5, 1.0 and 1.5 in shaking incubator (37 °C 250 rpm). Cell cultures were harvested (3000 xg for 5 minutes) and resuspended with PLM to obtain the same cell count per mL (3*10^8^ CFU/mL). The cultures were then infected with lysate produced from ISA199 (*E. coli* C600 containing P1kc and 12 kbp phagemid). Infected cultures were plated on LB+Cm to count CFU/mL.

### Large phagemid infection rate experiments

PM-9 (9.7 kbp), PM-12 (12.8 kbp), PM-24 (24.0 kbp), and PM-42.0 (42.0 kbp) phagemids were transformed into *E. coli* C600 containing only P1kc:10kb or P1kc:10kb and aTc-MP. Phage lysate was produced from each strain as above and then used to infect wild type *E. coli* C600 or *E. coli* C600 containing P1kc::10kb. Infected strains were plated on LB+Cm plates to count CFU/mL.

### Infecting different *E. coli* Strains

The 12 kbp phagemid was transformed into *E. coli* C600, MG1655, and Nissle containing P1kc:10kb::KanR. 3 biological replicates from the transformed strains were grown overnight in LB+Cm and used for phage production. Phage lysate from each strain was used to infect wild type *E. coli* C600, MG1655, and Nissle.

### Flip recombinase to delete *KanR* from P1kcΔcoi::KanR and P1kc:10kb::KanR

Adapted from barricklab.org^43^. In brief, *E. coli C600* containing P1kcΔcoi::KanR (ISA137) and P1kc:10kb::KanR (ISA138) were first transformed with pCP20 and plated in LB+Ampicillin agar plates and grown overnight at 30 °C. Grown colonies were inoculated into 1.0 mL of LB in eppendorf microcentrifuge tubes and grown overnight at 43°C to induce FLP recombinase expression and plasmid loss. Colonies were then screened for loss of *KanR* from the phage genome via plating in LB+Kan, LB+Amp and LB only plates.

### Single and double Stop codon reversion

*AmpR* was inserted into a phagemid backbone (ISA012) via Gibson assembly. Mutations were introduced via Q5 SDM to introduce either one or two stop codons in *CmR*. This phagemid, with a difunctional *CmR,* was transformed into *E. coli* C600 containing aTc-MP and P1 phage (ISA308, ISA311 and ISA363). 3 biological replicates were grown overnight in LB containing 1% glucose, kanamycin, and ampicillin. The overnight cultures were plated in LB/Cm agar plates to check for escapers and background aTc-MP activity; no escapers were detected. The cultures were spun down at 4000 xg for 3 minutes in eppendorf microcentrifuge tubes and washed with 1X PBS twice to remove the residual glucose from the media. Washed cells were then transferred to LB media containing kanamycin and ampicillin (1:1000) and with or without 200 ng/mL aTc to induce aTc-MP. The cultures were plated after 8 hours or 16 hours of induction on LB+Cm plates to count the number of resistant colonies. Random colonies were picked to verify stop codon reversion.

### Isolation of recessive mutations in p15A-sfGFP via IDE (see Supplementary Methods)

The p15A origin and *sfGFP* were inserted into ISA012 via Gibson assembly (ISA010). The resulting phagemid (p15A-sfGFP) was transformed into *E. coli* C600 containing P1kc:10kb and aTc-MP (ISA384). In short, the p15A-sfGFP phagemid was mutated, transferred to fresh cells, and selected for higher fitness via outgrowth a total of 4 times. The phage lysates produced from each round’s stationary phase cultures were used to infect wild type *E. coli* C600. These strains were analyzed via flow cytometry to compare their fluorescence to that of empty vector, wild-type, and cell cultures that passed through the steps with the exception of mutagenesis. Flow cytometry data was analyzed with FlowJo (FlowJo LLC). The *sfGFP* gene from 5-10 colonies was amplified via PCR and sequenced by Sanger sequencing to detect mutations.

### Improving pSC101-sfGFP fluorescence via IDE (see Supplementary Methods)

The pSC101 origin and *sfGFP* were inserted into ISA012 via Gibson assembly. The resulting phagemid (pSC101-sfGFP) was then transformed into *E. coli* C600 containing aTc-MP and P1kc:10kb (ISA426). The resulting strain was mutated 2 times before selection for increased colony fluorescence. Colonies that exhibited higher GFP fluorescence were visually identified and analyzed via flow cytometry to quantify GFP production. Mutations that were found in pSC101 were introduced to the unevolved phagemid via Q5 SDM, transformed into *E. coli* C600, and analyzed via flow cytometry to confirm their causal effects.

### Tagatose evolution (see Supplementary Methods)

The tagatose pathway from *Bacillus licheniformis* (ATCC^®^ 14580™) was inserted into ISA012 via Gibson assembly. The phagemid was then transformed into *E. coli* C600 containing aTc-MP and P1kc:10kb. The resulting strain was evolved through two rounds of diversification and selection as shown in Figure S12. The resulting phage lysate was used to infect wild-type *E. coli* C600, and the resulting cells were plated on tagatose minimal media agar plates. Large colonies were picked and grown overnight in LB media containing chloramphenicol. The cultures were then washed with tagatose minimal media and grown for 40 hours in a plate reader (see methods: growth curves). The pathways in the colonies with faster growth rates were sequenced via Sanger sequencing. Resulting mutations were then reintroduced to unevolved phagemid via Q5 SDM and analyzed again via growth in tagatose minimal media.

### Tagatose Minimal Media

The selection media for the tagatose minimal media was adapted from Van der Heiden, *et al.* (2013)^37^ with few modifications for *E. coli* C600. 20x Salt solution was composed of KH_2_PO4 (54.4 g), K_2_HPO_4_ (208.8 g), and NH_4_Cl (12 g) for 1 L solution. 500x mineral solution was composed of MgCl_2_·6H_2_O (1 g), CaCl_2_·2H_2_O (0.25 g), FeCl_2_·4H_2_O (25 mg), ZnSO_4_·7H_2_O (25 mg), CoCl_2_·H_2_O (12.5 mg), CuSO4·5H_2_O (0.5 mg), and MnSO_4_·H_2_O (0.14 g) in 100 ml solution. 100x vitamin mix (50 ml) was composed of 5 mg of thiamine-HCl, nicotinic acid, folic acid, d-l-pantothenic acid, d-biotin, leucine, lysine, homoserine and riboflavin and 10 mg of pyridoxal-HCl. 0.1% (wt/vol) I-Casamino Acids and 1% (wt/vol) tagatose were added to the minimal media.

### Growth curves

3 biological replicates of each isolate were grown overnight in 96-deep-well plates (VWR International, cat #10755-248) in LB media. Saturated cultures in 96-deep-well were spun down at 2500 xg for 10 minutes, the supernatant was discarded, and the pellet was resuspended in tagatose minimal media. These suspensions were spun again at 2500 xg for 10 minutes, the supernatant was discarded, and the pellet was resuspended in tagatose minimal media one last time. This culture was then transferred to 96-well-plates (Costar, Corning™ 3788) containing tagatose minimal media (1:200 dilution) and grown for 40 hours in a plate reader (BioTek Synergy™ H1, Shake Mode: Double Orbital, Orbital Frequency: continuous shake 365 cpm, Interval: 10 minutes).

---

**Figure S1.**
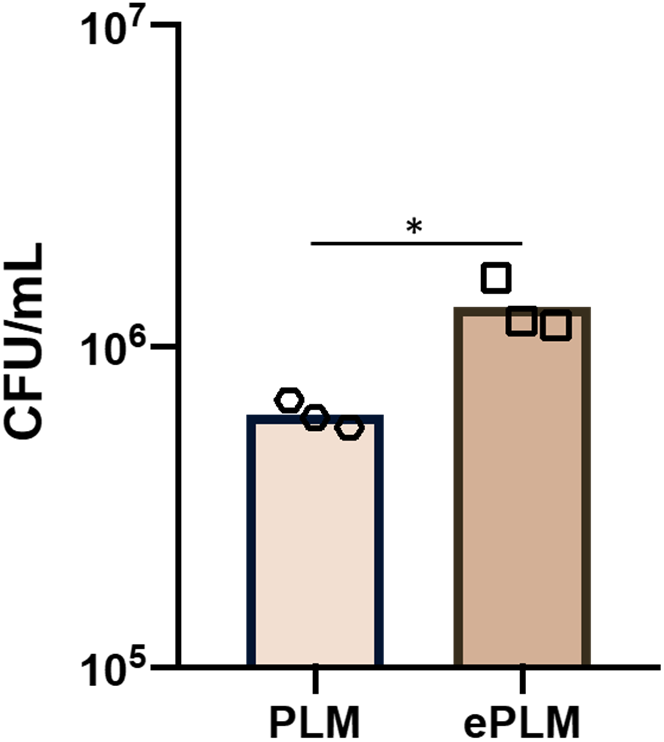
Increasing salt composition (MgCl_2_ and CaCl_2_) in the growth medium influences phage infection rate. Each dot represents one biological replicate.

**Figure S2.**
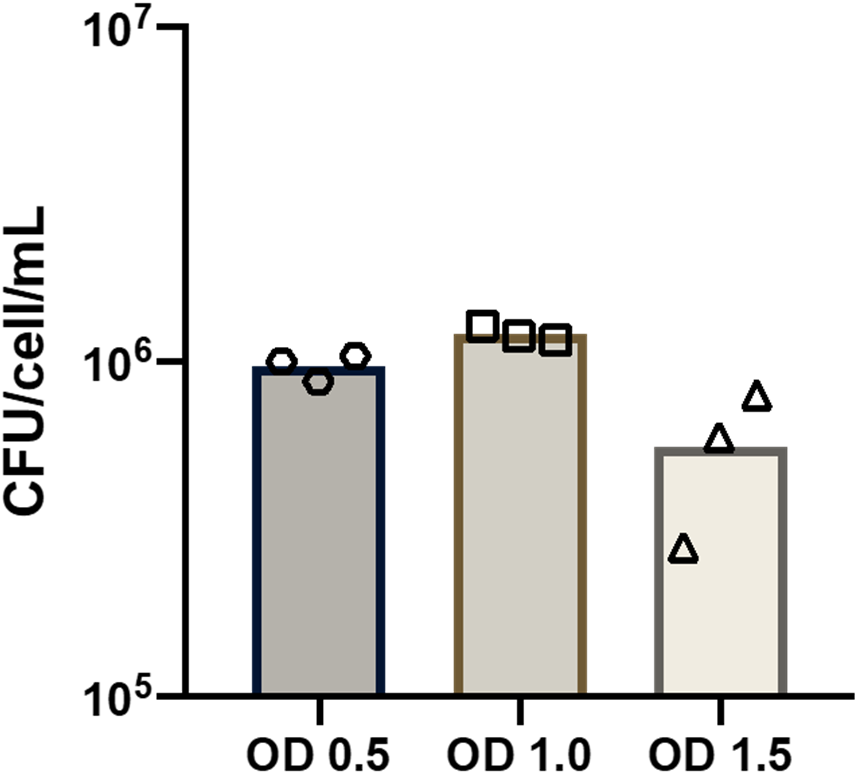
The physiological state of cells (*E. coli* C600) affects phagemid infection rate. Each dot represents one biological replicate.

**Figure S3.**
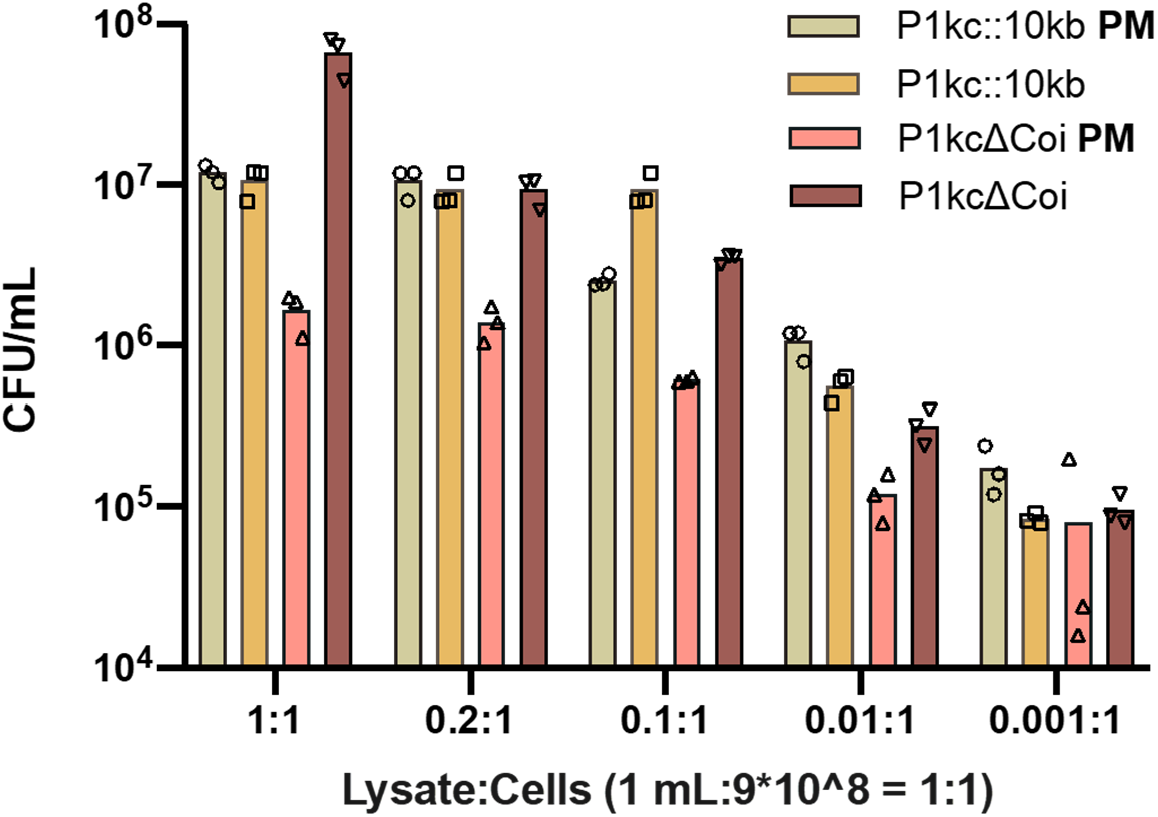
The ratio of lysate produced from strains containing phagemid and P1::10kb or P1ΔCoi affects the amount of the size of the library passaged in each cycle. Each dot represents one biological replicate.

**Figure S4.**
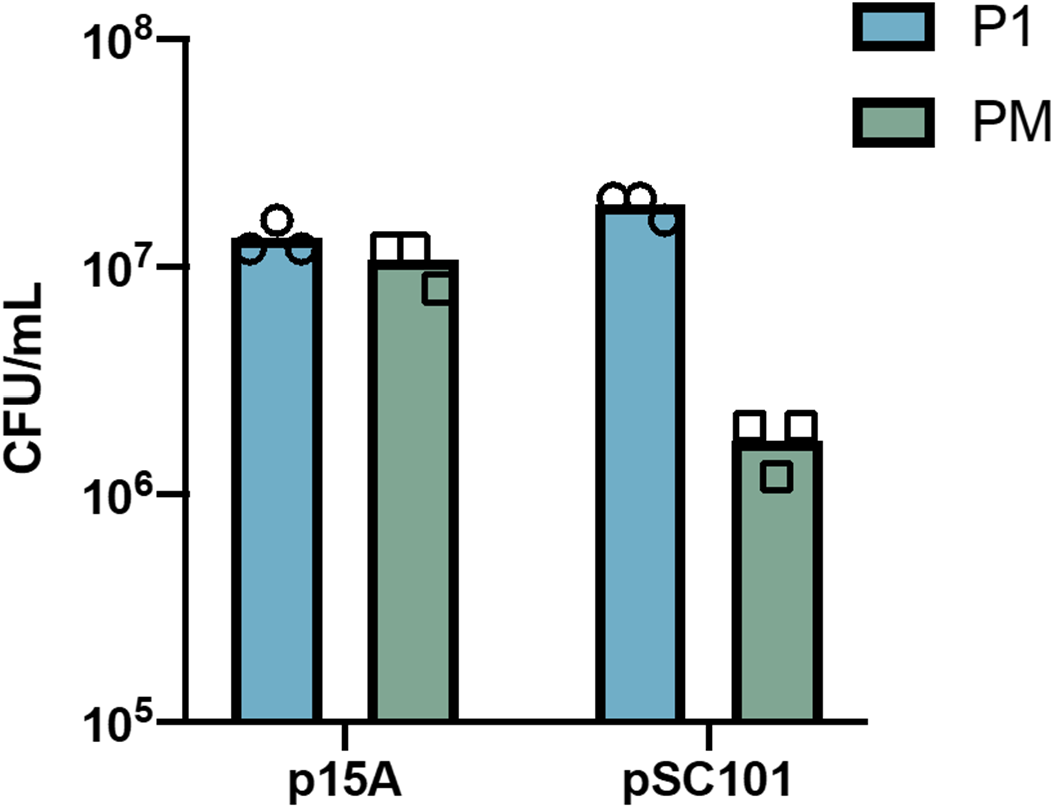
The copy number of phagemid impacts packaging and infection rates in the context of P1kc::10kb. Each dot represents one biological replicate.

**Figure S5.**
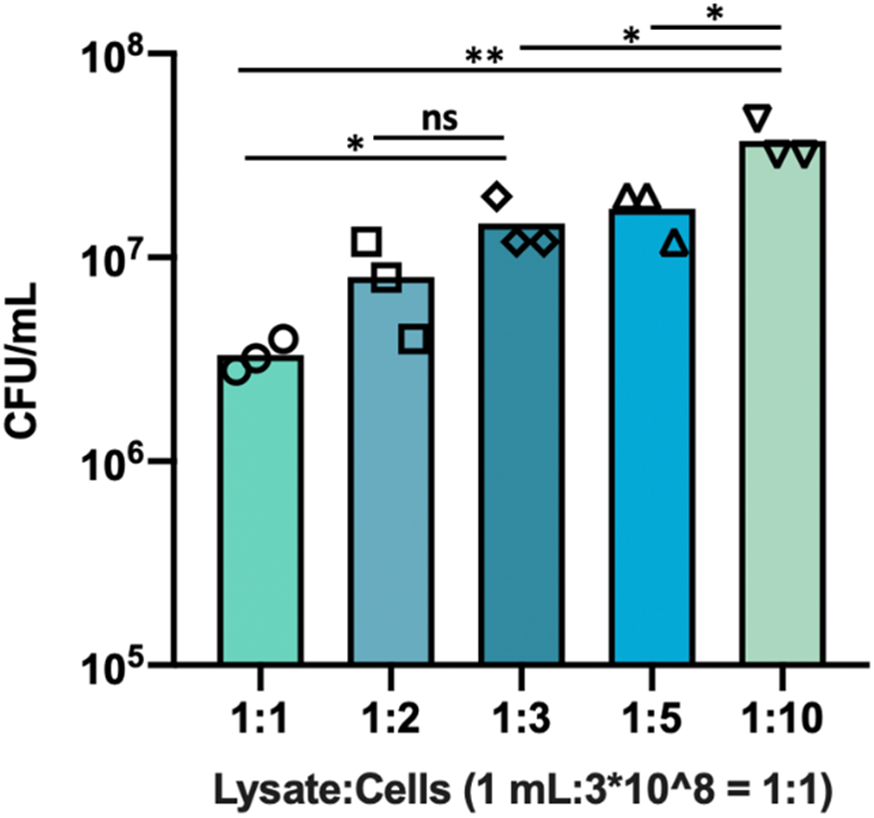
The amount of infected cells grown to OD 1 affect the size of the library passaged in each IDE cycle. 3 biological replicates of *E. coli* C600 cells were grown to OD 1 and concentrated to 1x, 2x, 3x, 5x and 10x and then infected with phage lysate produced from *E. coli* C600.

**Figure S6.**
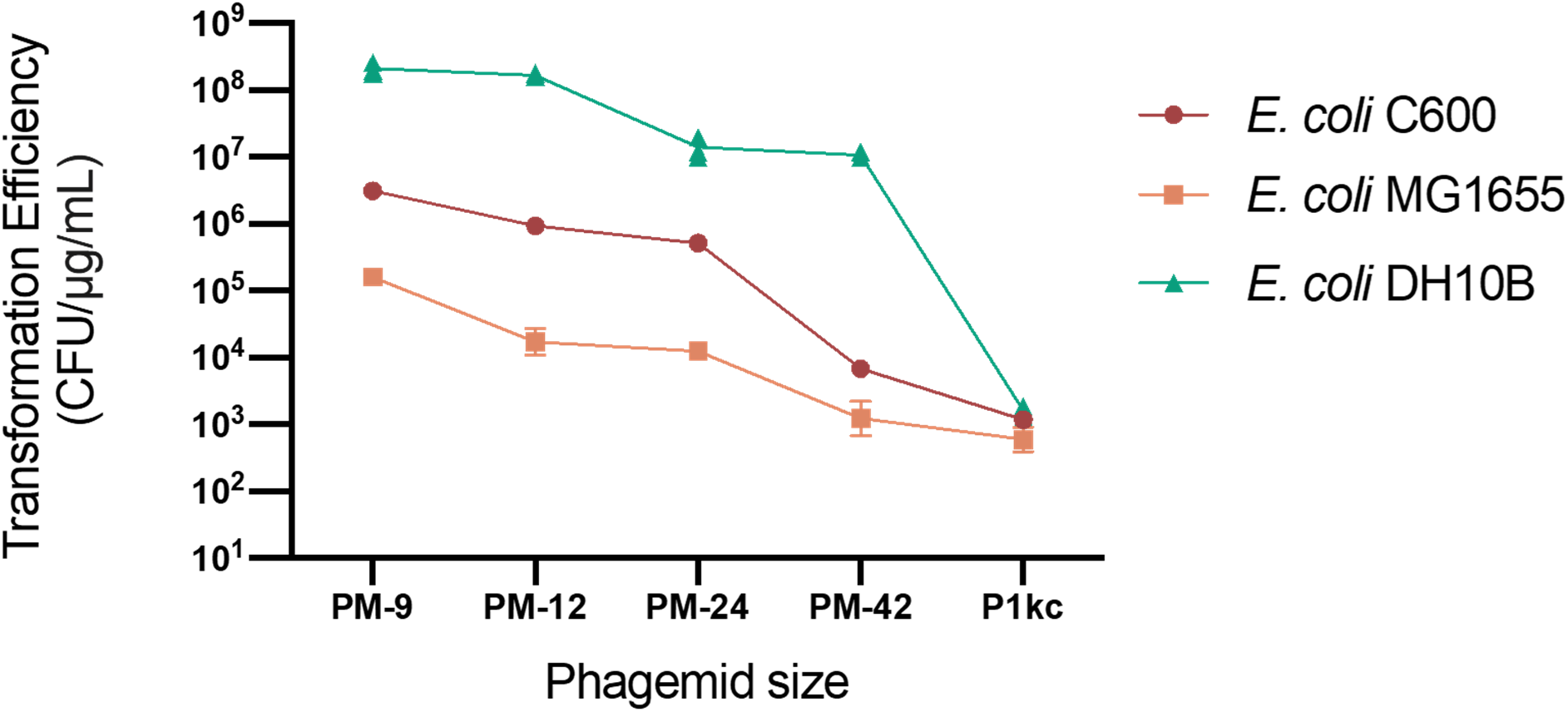
Transformation efficiency of *E. coli* C600, MG1655 and DH10β by different size phagemids. 1 μg of each phagemid was transformed into freshly prepared electrocompetent cells.

**Figure S7.**
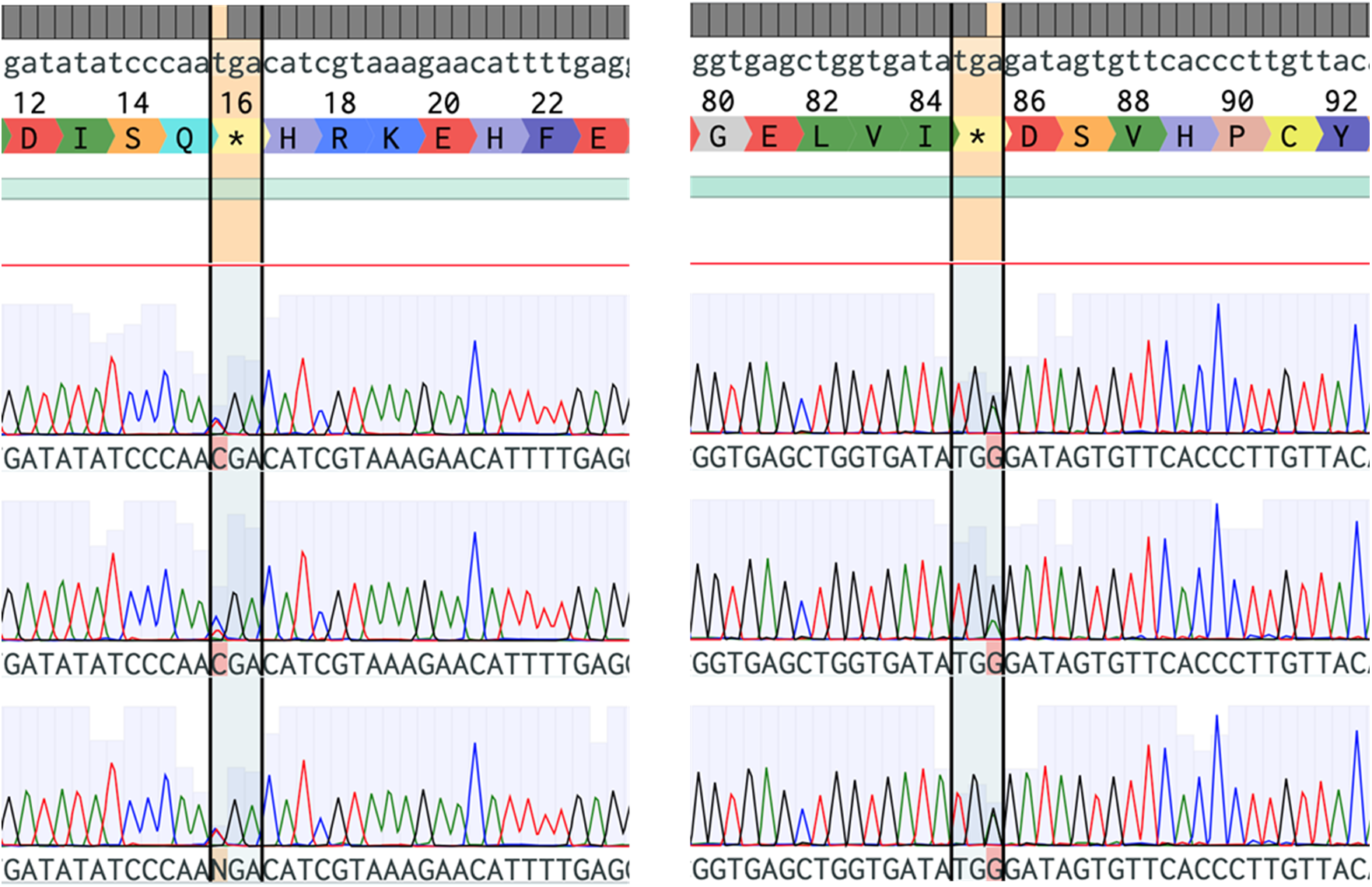
Sanger sequencing shows the reversion of two premature stop codons inserted in *CmR* at codons 16 and 85. Both codons were originally Trp (TGG). Double peaks indicate that the cells have two copies of the phagemids (one with stop codon and one with reverted stop codon).

**Figure S8.**
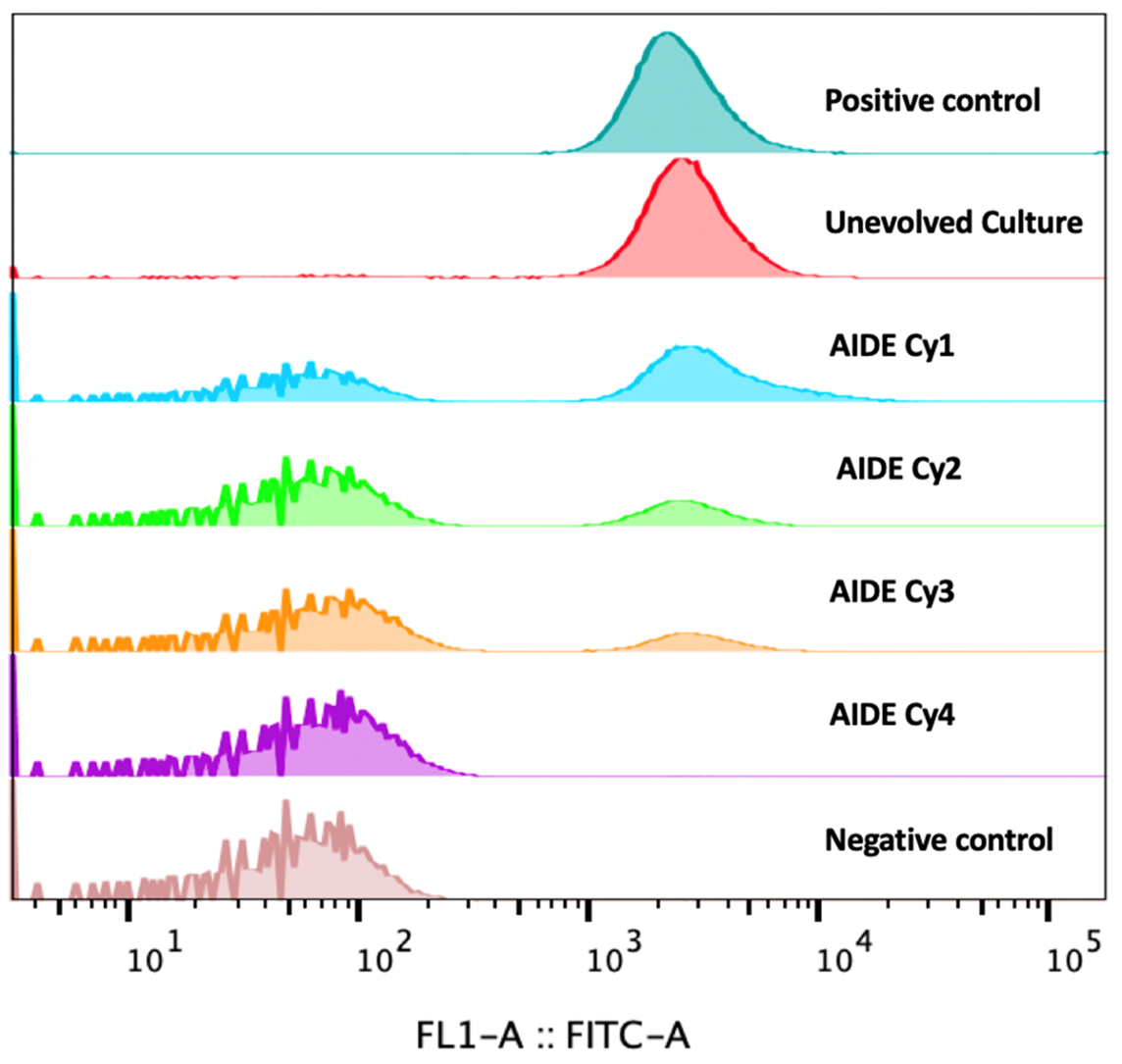
Directed Evolution of *sfGFP* using IDE. *sfGFP* phagemid was evolved to test the ability of IDE to passage recessive variants between cells across multiple rounds. The *sfGFP* phagemid went through 4 IDE cycles and was compared using flow cytometry to 1) a phagemid that was not mutagenized but that went through the rest of the IDE steps (unevolved culture), 2) negative controls (harboring an empty plasmid), and 3) positive controls (the starting plasmid).

**Figure S9.**
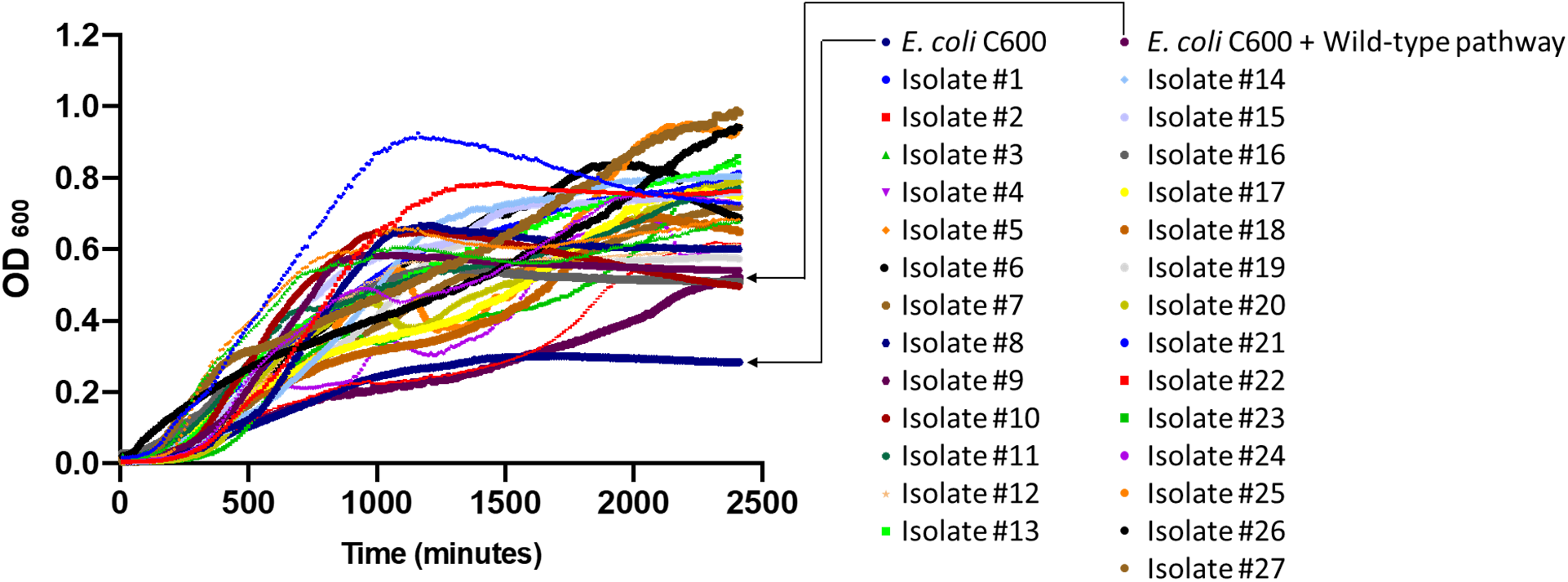
Growth curves of 27 tagatose variants. 27 colonies were picked from tagatose agar plates after 2 rounds of mutagenesis and selection. These colonies were grown in liquid tagatose media in a plate reader with wild-type *E. coli* C600 and *E. coli* C600 containing the wild-type pathway.

**Figure S10.**
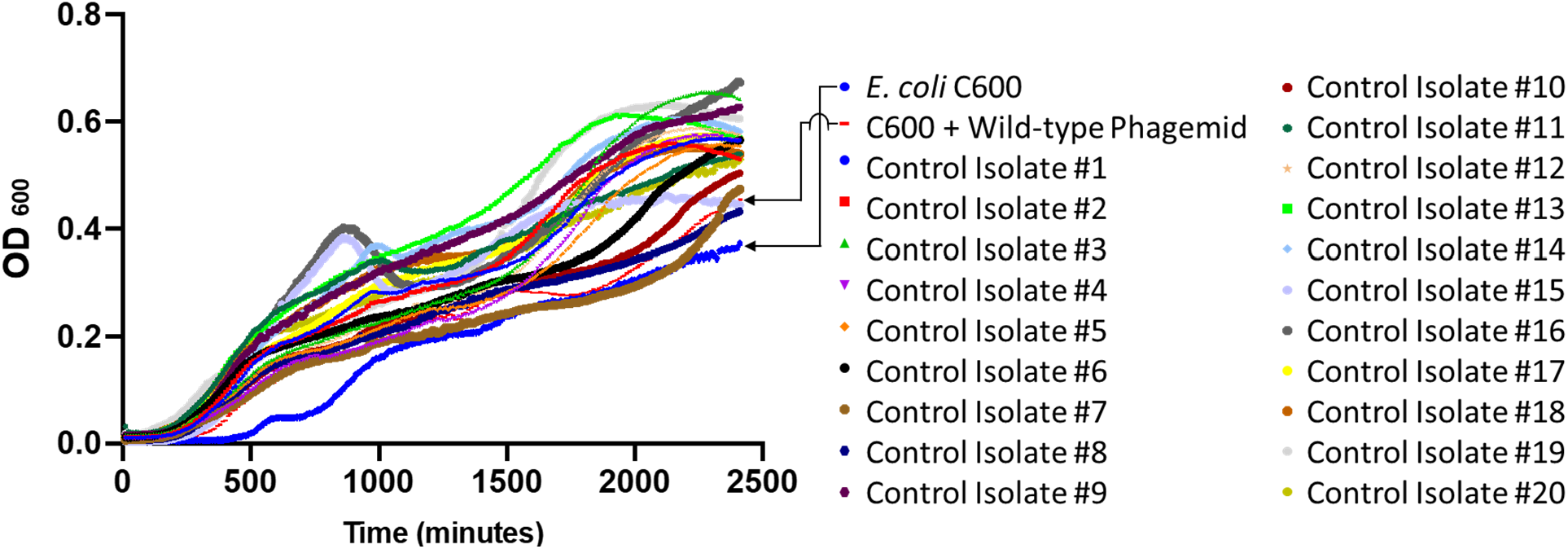
Growth curves of control colonies. 20 colonies were picked after 2 rounds of selection without mutagenesis. Colonies were picked from tagatose agar plates and grown in a plate reader.

**Figure S11.**
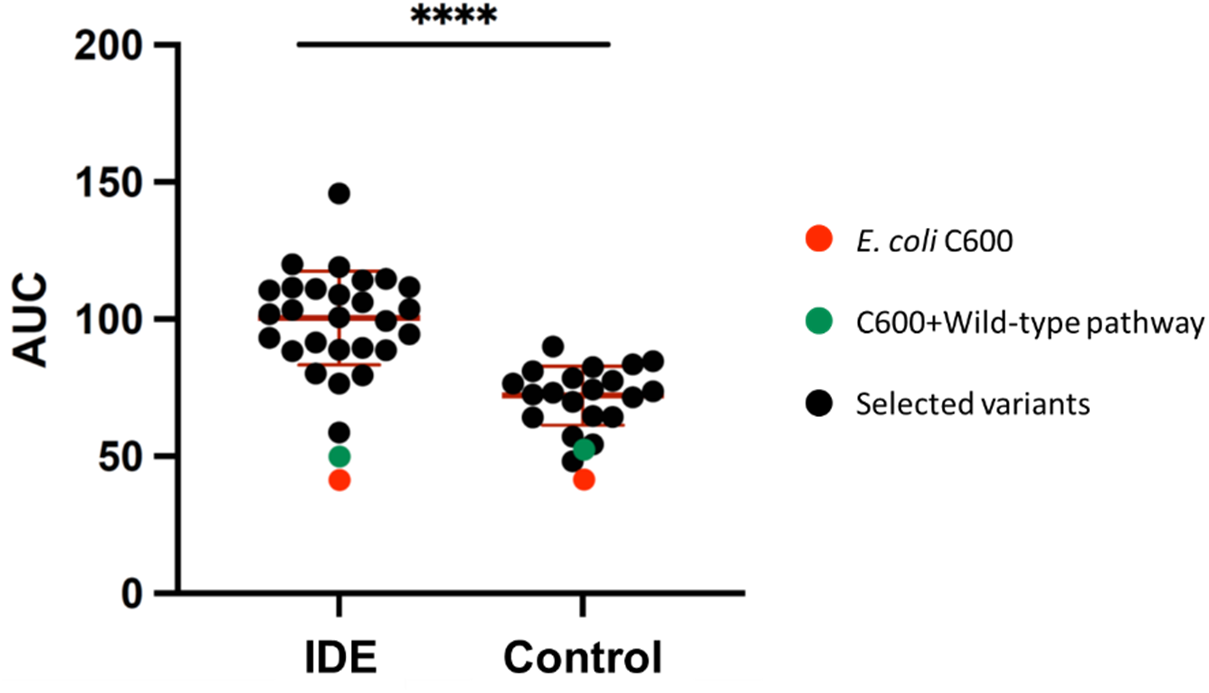
Comparing growth characteristics of variants isolated from IDE and Control tagatose selections. Variants from IDE and control selections were grown in tagatose minimal media and optical density was measured over time in a microplate reader. AUC (Area Under the Curve) is calculated by summing OD600 values obtained over the course of the experiment. **** - p<10^-8, Student’s T test.

**Figure S12.**
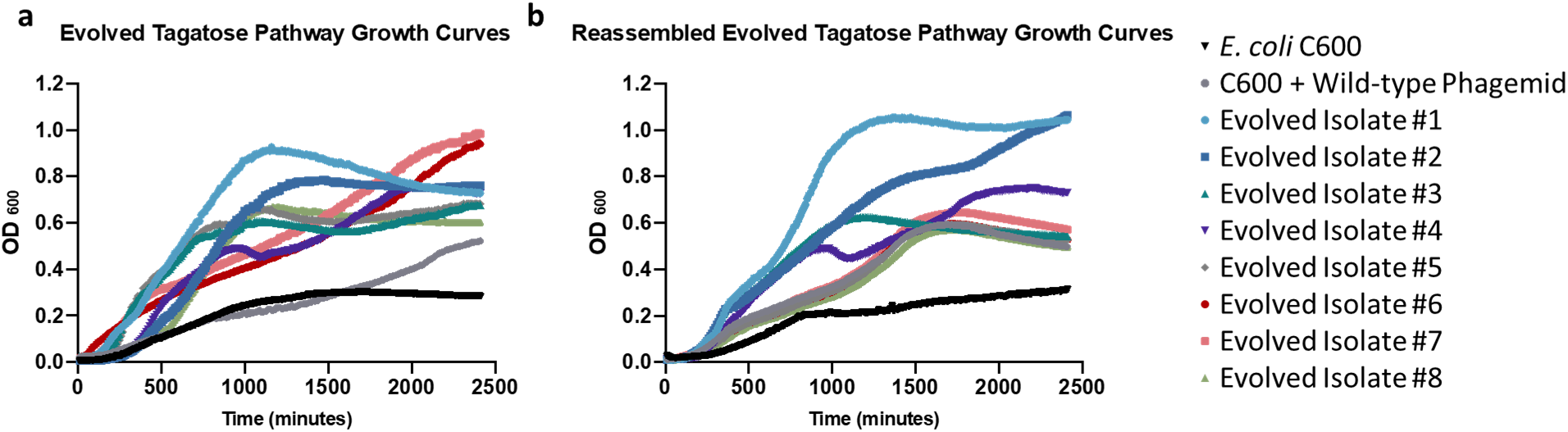
Growth curves of the selected tagatose variants. **a.** 8 variants were picked out of 27 isolates that were picked initially from tagatose agar plates and confirmed in plate reader. **b.** Tagatose pathways from the 8 selected variants were assembled into unmutated backbone and grown in tagatose minimal media.

**Table S1.**
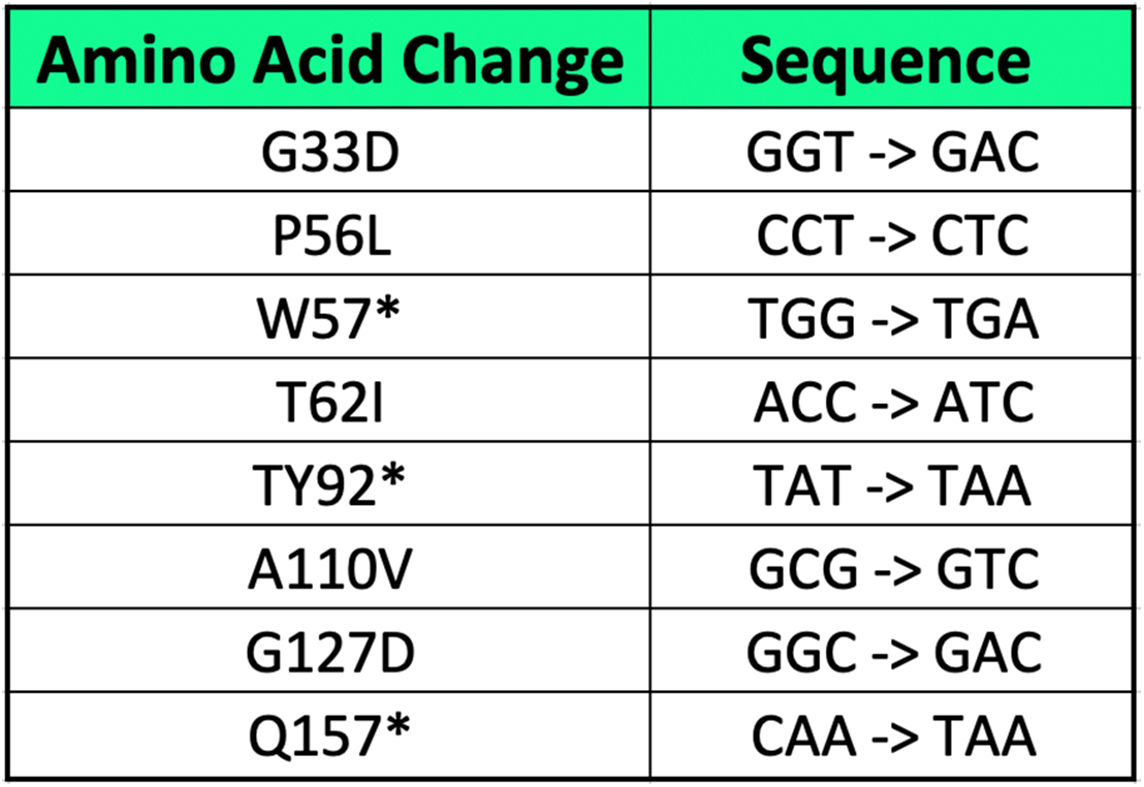
Detected mutations in the p15A-sfGFP phagemid after 4 IDE cycles. All mutations were detected in sfGFP. *=stop codon.

| Amino Acid Change | Sequence |
| --- | --- |
| G33D | GGT -> GAC |
| P56L | CCT -> CTC |
| W57* | TGG -> TGA |
| T62I | ACC -> ATC |
| TY92* | TAT -> TAA |
| A110V | GCG -> GTC |
| G127D | GGC -> GAC |
| Q157* | CAA -> TAA |

## Acknowledgements

We thank members of the Crook Lab, and Daphne Collias, Dr. Chase Beisel, Dr. Jennie Fagen, and Dr. Janetta Hakovirta for valuable discussions and input. We also thank the labs of Dr. Christopher Anderson (UC Berkeley) for phagemid constructs (Addgene #40782, #40783 and #40784), Dr. David R. Liu (Harvard University) for the MP6 plasmid (Addgene #69669), and Dr. Chase Beisel for wild type and engineered P1 bacteriophages. This work was supported by startup funds from North Carolina State University’s Chemical and Biomolecular Engineering (CBE) Department and NCSU’s Faculty Research and Professional Development Fund. I.S.A. is supported by NCSU CBE startup funds and the Ministry of Higher Education - Oman.

## Author contributions

I.S.A. and N.C. designed and conceived the study. I.S.A. and D.J.H. conducted all experiments. N.C. supervised the research. I.S.A., D.J.H., and N.C. wrote the manuscript.

